# Global change affects differently at local scales animal populations: why Starlings still expands to the south while other temperate species go to the north?

**DOI:** 10.1101/2021.08.05.455352

**Authors:** Alexandra Rodriguez, Giuseppe La Gioia, Patricia Le Quilliec, Damien Fourcy, Philippe Clergeau

## Abstract

Global change, which regroups global warming, landscape transformations and other anthropic modifications of ecosystems, has effects on populations and communities and produces modifications in the expansion area of species. While some species disappear, other ones are beneficiated by the new conditions and some of them evolve in new adapted forms or leave their ancient distribution area.

As climate change tends to increase the temperature in several regions of the world, some species have been seen to leave areas in equatorial regions in order to join colder areas either towards the north of the northern hemisphere or towards the south of the southern one.

Many birds as have moved geographically in direction to the poles and in many cases they have anticipated their laying dates. Actually, two tit species that use to lay their eggs in a period that their fledging dates synchronize with the emerging dates of caterpillars are now evolving to reproductive in periods earlier than before the climate change. Several species are reacting like that and other ones are moving to the north in Europe for example. Nevertheless, and very curiously, European starling, *Sturnus vulgaris*, populations are behaving on the contrary: their laying dates are moving towards later spring and their distribution area is moving towards the south. In this study we explore and discuss about different factors that may explain this difference from other birds.

## Introduction

Climate change is divided into six phenomena : modifications in CO2 concentrations, incesasing temperatures, increasing ocean level, melting ice, increasing of desert areas and multiplying extreme events as heat waves, floods, storms and cyclones. These modifications are the result of planet anthropisation and have several consequences on organisms’ life history from local to global scales (Parmesan 2006).

When climate warms vetebrate have three solutions : to move towards colder areas, to develop local adaptations or to disapear. The most common mechanism is habitat tracking (Raia et al. 2012) which is a mechanism sometimes invoked by scientists to explain morphological stasis. In this case populations can simply follow their preferred habitat rather than adapting to new conditions in their original distribution area. In the case of birds, they follow the thermic envelop to which they are already adapted by moving to higher latitudes or altitudes.

Several studies are conducted on bird’s vulnerability to climate change are they are expected to be particularly touched by this pheneomenom (Pearce-Higgings et al. 2014). Other authors suggest that an increase in 1°C of the temperature could lead to the extintion of 100 to 500 species.

Several life history traits are modified with diferent intensities as a response to climate change. For exemple, birds tend to nest earlier in spring (Crick et al. 1997). These changes in life history traits can be due to the phenotipical plasticity when an individual adjusts its response to an environmental signal as immediate temperature : birds lay eggs later when spring is cold and earlier when its warm (Thomas et al. 2001). Birds use proximal signals as photoperiod and temperature, to avoid gaps between the dates of laying eggs and caterpillars hatching that depend themselves on buds break dates (Lambretchs et al. 1997). In Great tits, Parus major, it has been observed that ringed individuals presented a laying date that was in advance of 14 days from the habitual date which allows them to feed their chicks during the presence of caterpillars (Chamantier et al. 2008). In the same way, it has been demonstrated that it is possible to modify the migratory behavior of birds as *Sylvia atricapilla* by applying selective presures in aviaries. A little population which was partialy migratory at the start of the experiment can elvolve by artificial selection into a totally migratory population after three generations or a totally sendentary population after six generations (Berthold et al. 1991).

It has been reported that several species from the most studied terrestrial groups have presented a shift in their distribution as a majority of birds and buterfly species distributions have moved towards higher latitudes (Devictor et al. 2008, Devictor et al. 2012). This phenomenom is remarquable from the nineties in 200 of the 500 nesting European species (Burton 1995) and it has been accelarated from 2000. However, even if climate warming has an important impact on the observed and measured shifts it is very important to make the distinction between climate change and other environmental factors as habitat structure and biological species adaptations to novel conditions. In the northern hemispher birds actually move towards the north in response to the climate change but they do it slower than predicted as the birds travelled distance and the distance between two isotherms is about 182 km (Devictor et al. 2007). Moreover, it has been observed that species like European starling, Sturnus vulgaris, do not move to the northern regions of Europe but they move towards southern Europe (Ferer et al. 1991).

This species native from central and northern Europe (Feare 1984) was introduced in several regions of the world. In XIX ° century, it has been introduced in the United States. In 1899 it has been introduced in South Africa (Cooper and Underhill 1991) and in 1882 in Australia (Long 1981). More recently, in 1987 (the last reported introduction) the species has been brought to Argentina (Peris et al. 2005). In the regions of the southern hemisphere (South Africa and Argentina) the species is expanding from the south to the north (Peris et al. 2005, Cooper and Underhill 1991) whereas in the northern hemisphere (in Europe and in North America) the species has been expanding to the south reaching southern Mexico in the American continent (Rodríguez-Estrella 1997) and the north of Spain and the south of Italy and Greece in the European continent (Heldbjerg et al. 2019). It has been even reported that English and Finish populations are in decrease, which let us doubt between and invasive process in the south or a translation of the expansion area of the species (Solonen et al. 1991, Rintala et al. 2003).

Thirty years of observation of the species in Brittany region in the northwestern France indicate that Starlings contrary to other studied species are not anticipating their laying dates but they are getting them latter (Clergeau personal observations). Fourty years ago, starlings of this region laid eggs by the first april, now they have been doing it by 12^th^ april. Starling thus appear as an interesting species to enlarge studies on effects of global change as it is like a counterexample that let us think that many aspects of effects of global change are not being took into account or that there are several factors that are acting on populations changes.

In this study, we focused on Italian species’ invasion and tried first to reconstruct and confirm species’ distribution in the southern regions of the country. Then we tested the hypothesis that the habitats that have not been colonized before by the species in this region became suitable and invasible by recent anthropic changes in climate and landscape characteristics.

We explore here some factors that can be responsible of modifications observed in starling populations in southern Europe and suggest some explanative hypothesis to it recent switch to the south of the continent.

## Material and methods

### 1. Confirming Starling distribution and reconstructing it’s history in Italy

#### Literature approach

We used different European Atlas (French and Italian) with presence-absence maps to define the areas of presence of the starling and local Italian ornithological journals in order to take into account the field observations by the local ornithologists.

We verify on the most recent Italian maps, which are the suitable regions for the species in that country.

#### Field observations and confirmation

We carried on precise field observations in Apuglia and Calabria region in order to identify the breeding colonies of the propagation front (confirm those reported by Italian ornithologists and detect if they were new ones).

Precise observations in Apuglia region were conducted at the same time as we displayed behavioural studies on populations of the region during breeding periods of spring 2007 and 2008 (march and april).

We realised transects in Apuglia and Calabria region by car following regional routes and noted each time that a breeding colony was identified.

### 2. Testing hypothesis of possible causes of Starlings’ move to the south

#### Eliminating the hypothesis of a lack of cavity breeding sites

As Eurpean starling, *Sturnus vulgaris*, is a cavernicole species that makes it nests in tree holes or in holes in buildings and houses or under the roofs at more than five meters from the ground. Cavity availability combining both the presence of the hole and its’ height can be a limiting factor to the settlement of starlings’ populations. In areas with high densities of starlings as in Britany it has been reported that fights between couples took place into the nests boxes (Jean-Pierre Richard personal communication) even when cavities where free but when only some of them was interestingly located (presenting both shadow and sun).

Apuglia region is located on the southern east coast of Italy along the Adriatic sea. In this region mainly of the starling rural populations’ breed on the cliffs. We hypothesized that there were enough cavities to nest thus that they will not be a limiting factor for starling settlement.

The Apuglia urban populations construct their nests in holes in the top of the big recent buildings typical of southern Italy architecture or in the road lamps. As there are few old trees with cavities and as human disturbance can be more or less important in the Italian roof that are often associated to terraces with possibility of human presence we have doubts about the impact of cavity availability in this habitat.

In order to test the hypothesis that cavities availability was not a limiting factor for starling expansion both in rural and urban areas we put 30 nest boxes in different areas of the southern propagation front.

##### Location

In all the cases, the nest boxes were put in areas actively frequented by starlings (near foraging grounds or immediately besides the breeding areas). We put nest boxes in the following cities and rural suroundings: Bari, Mola di Bari, San Cataldo, San Andrea and Otranto.

Characteristics of the nest boxes: the nest boxes were typical boxes for starlings and well accepted by the species in other European regions (green wood walls 25×17×17cm) (Mennechez and Clergeau 2006)

## Results

### 1. Starlings’ history in Italy

We confirmed that several of the sites where starlings were reported in Apuglia and Calabria where occupied by the species.

We confirmed the sustainability of starling populations in Apuglia region.

We confirmed the breeding area of Bari and Mola di Bari cities (central parks and University parks).

We observed ten individuals in the industrial area of Brindisi.

#### Data of verified sites

In San Cataldo we observed:

18 couples in San Cataldo breeding on nine street lamps (two adjacent nests by lamp)

30 feeding individuals at Specchia dell’Alto

20 feeding individuals at the woodland street.

Cliff between Rocca Vecchia and San Andrea:

250-300 individuals were observed

In rural Otranto:

Four couples were observed at the extreme edge of the continent: two couples at Torre del Serpe and 2 couples at the Faro.

#### We found tree new places where starlings have not been reported

- Urban Otranto: 1 couple nesting in the Castle
- Suburban Frigole: Five individuals eating near the Idrofor Tour
- One couple nesting in a lamp at suburban Frigole

#### We did not observe individuals in an area where they were reported

One year before (in 2006) Giussepe La Gioia had observed breeding couples in the area of Galiano. We did not find them any more in spring 2007 and 2008.

Taking into account the information of starlings’ presence in Italy depending on our different sources, we retraced the following chronology of starlings’ expansion (Figure 1 and 2).

**Figure 1.**
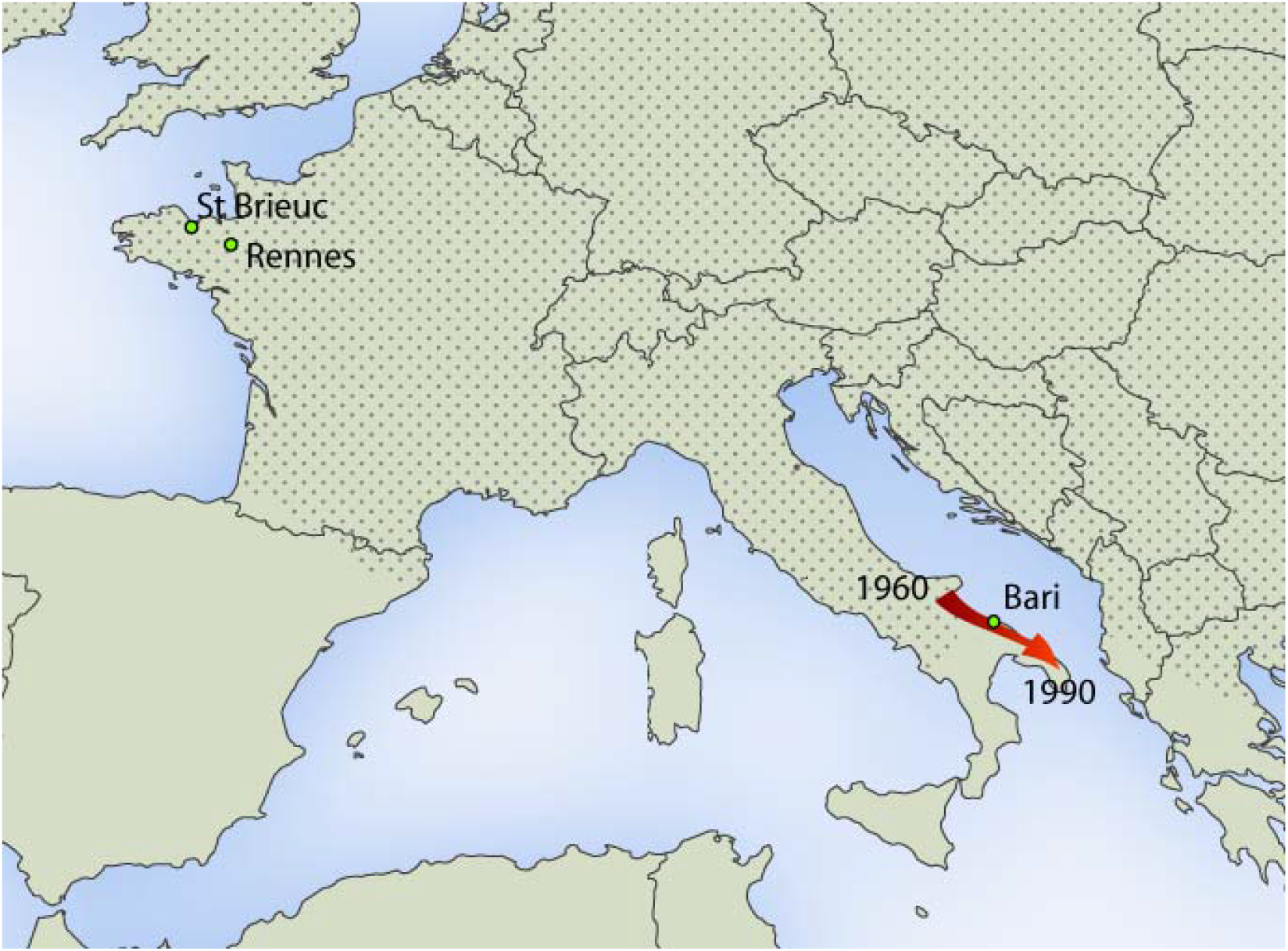
European starling expansion towards the south of Italy.

**Figure 2.**
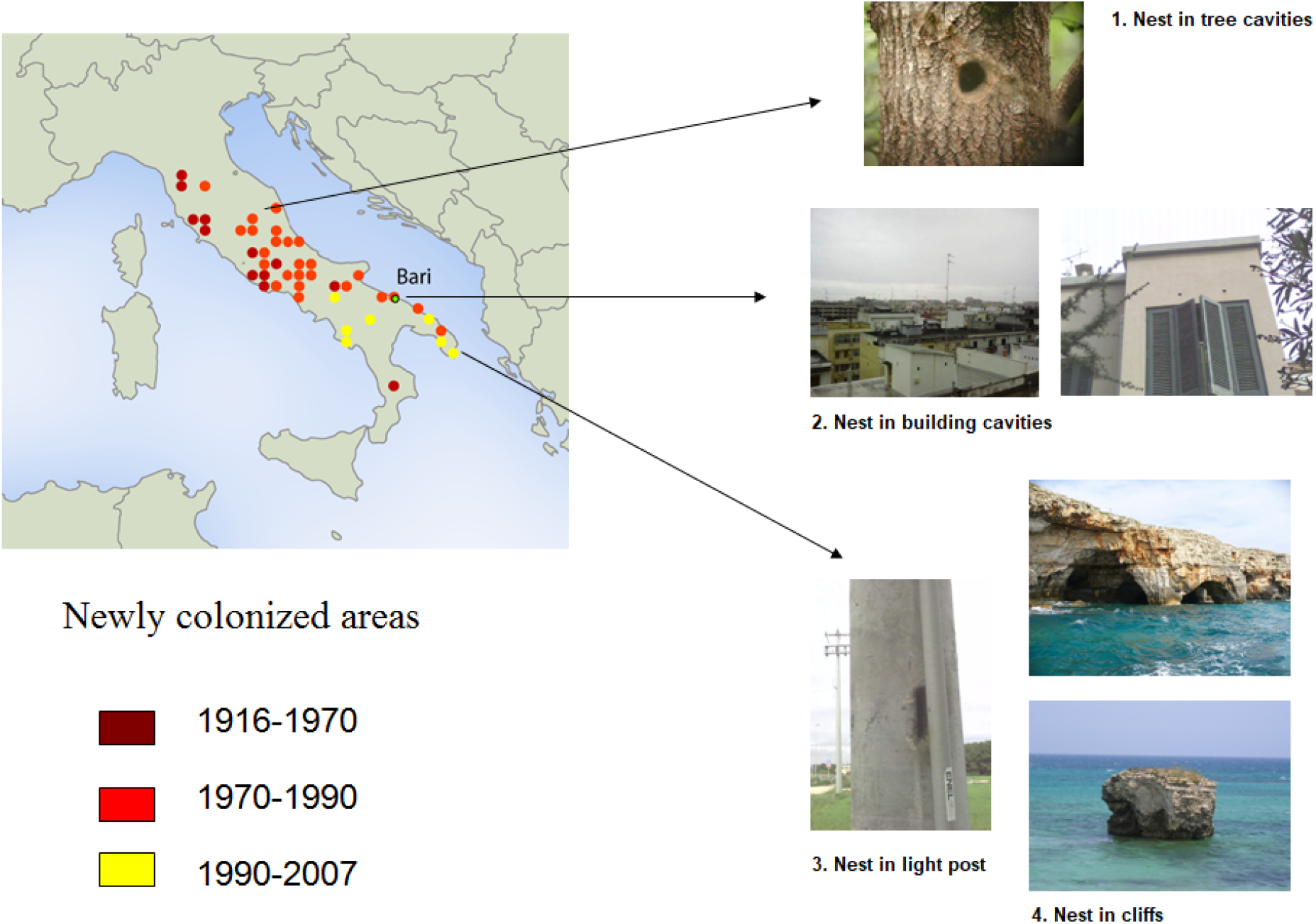
Dates of European starling settlement in Southern Italy and cavity nests.

As we can see in the figures, there is not a clear propagation front; the expansion seems to be done by little steps and radiation around primary foyers.

We can also see that there was a two-coast expansion. Starling seems to have been blocked during a period by Alpes Mountains as it is frequently the case with this species that does not occupy high areas. Then there was a rapid colonization of the north of Italy probably by birds arriving by the valleys from Swiss.

These first settlements stopped at the north of Italy. Density probably augmented and then they went to southern Italy.

They arrived by the 60’s to the north of Apuglia occupying cities like Bari, Mola di Bari and Brindisi.

Finally, they settled in the southern part of Apulia from the 70’s.

### 2. Eliminating the hypothesis concerning cavities

None of our 30 nest boxes was occupied by starlings. When we monitored them one year after they have been put on the buildings and trees, we have never seen starlings visiting them. We can thus think that there is not a lack of cavities or a demographic pressure in this area. Nevertheless, we have observed tits that have constructed their nests and that have chicks in our boxes. Usually when cavities are limiting factors for starlings they fight between them or with other species in order to get the good cavities (Mazgajski 2000). They usually win in confrontations with other species especially if they are smaller than starlings. As we have seen tits that have settled in the boxes we can think that there is not competition between this species and the European starling: they are probably well settled in the cliffs and for the moment they don’t seem to have problems with cavities availability. It has already has been reported that starlings used cliffs to nest in other regions of the world (Edwin 1971)

## Discussion

In this study, we confirmed that European starling occupies now rural and urban areas in southern Italy. By this way, we can thus confirm that the species distribution area has moved to the southern region of Europe, as they were no European starlings in the region before 1990. These observations are not in accordance with observations on other species of birds and even on species of butterflies that have moved to the northern regions of Europe towards higher latitudes, which can be explained in this case by tracking habitat phenomenon in the context of climate change (Devictor et al. 2008, Devictor et al. 2012).

In our study, we observed that nesting sites were not a limiting factor as the nesting boxes were not occupied by the starlings. We can think that there is enough availability of nesting sites in the region an that this factor may facilitate the settling of the European starling in this southern region of Europe. Other studies have shown that Sturnus vulgaris has biological and behavioural facilities to expand and colonize new habitats. In paticular it’s behavioral flexibility and social communication enhances colonisation processes. Actually, starlings relay on social cues and public information contained in chorus and visual information in order to locate suitable sites to feed and breed. This phenomenom is more relevant in colonization fronts, in particular in the Apuglia region of Italy where we observed higher proportions of starlings reacting to social information produced by decoys and playbacks (Rodriguez et al 2010, Rodriguez et al 2021a). Moreover, it has been shown that groups of starlings contain bold and shy individuals and that bold ones explore quickly movel habitats, novel food and novel objects and that they facilitate the colonization of new habitats (Rodriguez et al. 2021b). Finally, cognitive processes of habituation and learning allow the acceptance of new objetcs by the starlings in the suroundings of their feeding and breeding sites (Rodriguez et al. 2021c). This kind of tolerance towards novelty which corresponds to a behavioral flexibility may also explain the fact that starlings have moved towards new areas in southern Italy. In the region we studied, we reported two novel food items used by the starlings. Ten nests were observed containing starling parents feeding their chicks both with olives and dates which were entirely swallowed by the young birds (Alexandra Rodriguez personnal observations). Flexibility in the use of novel food items of warmer southern regions of Europe and of Northern Africa may be another factor explaining the facilities of the species to occupy this region.

Finally, we observed that southern Italy landscape has deeply changed during the past decades. Several touristic sites have developped along the Adriatic sea that offer sports facilities as tennis courts and golf fields. These kind of grounds which represent short grassland with feeding possibilities for the European starling have probably enhanced the spread of the species towards the southern region of Europe instead of the northern regions where there are not so many summer camps for touristic activities. More studies with quatitative data on lansdcape cover should be conducted to confirm or invalidate our hipotesis. However, at this stage we can conclude that European starling has several facilities to occupy the sourthern Europe as nests availability, behavioral flexibility and favorable feeding habitats that can explain its spread to the south and that higlights the fact that climate warming is not always the factor that drives species’ distribution shifts. In several cases perturbed habitats by anthropic activities may facilitate birds settlement (Marchesi 2002).

## Bibliography

Berthold, P. 1991. Genetic control of migratory behaviour in birds. Trends Ecol. Evol. 6: 254–257.

Burton, J. F. 1995. Birds and Climate Change. London, Christopher Helm.

Charmantier, A., McCleery, R. H., Cole, L. R., Perrins, C. M., Kruuk, L. E. B. & Sheldon, B. C. 2008. Adaptive phenotypic plasticity in response to climate change in a wild bird population. Science 320: 800–803.

Cooper, J., Underhill, L.G. 1991. Breeding, mass and primary moult of European Starlings Sturnus vulgaris at Dassen Island, South Africa. Ostrich 62: 1–7.

Crick, H. Q. P., Dudley, C., Glue, D. E. & Thomson, D. L. 1997. UK birds are laying eggs earlier. Nature 338: 526.

Devictor, V., Julliard, R., Couvet, D. & Jiguet, F. (2007) French birds lag behind climate warming. Nature Proceedings 10 p. DOI: 10.1038/npre.2007.1275.

Devictor, V., Julliard, R., Couvet, D. & Jiguet, F. 2008. Birds are tracking climate warming but not fast enough. Proc. R. Soc. Lond. B 275: 2743–2748.

Devictor, V., van Swaay, C. Brereton, T. Brotons, L. Chamberlain, D. Heliölä, J. Herrando, S. Julliard, R. Kuussaari, M. Lindström, Å. Reif, J. Roy, D.B. Schweiger, O. Settele, J. Stefanescu, C. Van Strien, A. Van Turnhout, C. Vermouzek, Z. WallisDeVries, M. Wynhoff, I. & Jiguet, F. 2012. Differences in the climatic debts of birds and butterflies at a continental scale. Nature Climate Change. 2, 121–124.

Feare, C. 1984. The Starling. Oxford University Press.

Ferrer, Xavier; Motis, Anna; Peris, Salvador J (1991). “Changes in the breeding range of starlings in the Iberian peninsula during the last 30 years: competition as a limiting factor”. Journal of Biogeography. 18 (6): 631–636

Heldbjerg, H., Fox, A. D., Lehikoinen, A., Sunde, P., Aunins, A., Balmer, D. E., Calvi, G., Chodkiewicz, T., Chylarecki, P., Escandell, V., Foppen, R., Gamero, A., Hristov, I., Husby, M., Jiguet, F., Kmecl, P., Kalas, J. A., Lewis, L. J., Lindstrom, A., … Weiserbs, A. (2019). Contrasting population trends of Common Starlings (Sturnus vulgaris) across Europe. Ornis Fennica, 96(4), 153–168.

Lambrechts, M. M., Blondel, J., Maistre, M. & Perret, P. 1997. A single response mechanism is responsible for evolutionary adaptive variation in a bird’s laying date. Proc. Natl. Acad. Sci. USA. 94: 5153–5155.

Long, J.L. 1981. Introduced Birds of the World. Universe Books, Agricultural Protection Board of Western Australia. pp. 21–493 New York. 330 p.

Mazgajski T. D. 2000. Competition for nest sites between Starling Stumus vulgaris and other cavity nesters — study in forest park. Acta om. 35:103–107

Marchesi, L; Sergio, F; Pedrini, P (2002). “Costs and benefits of breeding in human□altered landscapes for the eagle owl *Bubo bubo*”. Ibis. 144 (4): E164–E177

Mennechez, G. & Clergeau, P. 2006. Effect of urbanisation on habitat generalists: starlings not so flexible? Acta Oecologica 30: 182–191.

Michael, Edwin D (1971). “Starlings nesting in rocky cliffs”. Bird-Banding. 42 (2): 123.

Parmesan, C. 2006. Ecological and evolutionary responses to recent climate change. Annu. Rev. Ecol. Syst 37: 637–669.

Pearce-Higgins, J. W. & Green, R. 2014. Birds and climate change. Impacts and conservation responses. Cambridge, Cambridge Univ. Press.

Peris, S., Soave, G., Camperi, A.n Darrieu, C., Aramburu, R. 2005. Range expansion of the Eropean starling *Sturnus vulgaris* in Argentina. Expansion del Estornino pinto *Sturnus vulvaris* en Argentina. Ardeola 52: 359–364.

Raia P, Passaro F, Fulgione D, Carotenuto F. Habitat tracking, stasis and survival in Neogene large mammals. 2012. Biol Lett.. 8(1):64–6.

Rodríguez A, Clergeau P, Hausberger M (2010) Flexibility in the use of social information in the successful Sturnus vulgaris, experiments with decoys in different populations. Anim Behav 80: 965–973

Rodriguez A, Hausberger M, Henry L, Clergeau P. 2021a. Social information and biological invasions: chorus songs are more attractive to European starlings in more recently established populations. https://www.biorxiv.org/content/10.1101/2021.07.24.453665v1

Rodriguez A, Hausberger M, Henry L, Le Quilleq Clergeau P. 2021b. Population colonization history influences behavioral responses of European starlings in personality tests. https://www.biorxiv.org/content/10.1101/2021.07.24.453662v1

Rodriguez A, Hausberger M, Henry L, Le Quilleq Clergeau P. 2021C. Habituation, task solving and memorization may facilitate biological invasions: the Starling example. https://www.biorxiv.org/content/10.1101/2021.07.24.453664v1

Rodríguez-Estrella. 1997. European starlings nesting in Southern Baja California, Mexico. Wilson Bulletin 109: 532–535.

Rintala, J., Tiainen, J., Pakkala T. 2003. Population trends of the Finnish starling Sturnus vulgaris, 1952-1998, as inferred from annual ringing totals. Ann. Zool. Fennici. 40: 365–385

Rollins LA, Woolnough AP, Wilton AN, Sinclair R, Sherwin WB. (2009). Invasive species can’t cover their tracks: using microsatellites to assist management of starling (Sturnus vulgaris) populations in Western Australia. Molecular Ecology, 18, 1560–1573.

Rollins LA, Woolnough AP, Sinclair R, Mooney NJ, Sherwin WB. (2011). Mitochondrial DNA offers unique insights into invasion history of the common starling. Molecular Ecology, 20, 2307–2317

Sekercioglu, C. H., Schneider, S. H., Fay, J. P. & Loarie, S. R. 2008. Climate change, elevational range shifts, and bird extinctions. Cons. Biol. 22: 140–150.

Solonen, T., Tiainen, J., Korpimäki E., Saurola P. 1991. Dynamics of Finnish Starling Sturnus vulgaris populations in recent decades. Orinis Fecnica 68: 158–169

Thomas D. W., Blondel J., Perret P., Lambrechts M. M. & Speakman J. R. 2001. Energetic and fitness costs of mismatching resource supply and demand in seasonally breeding birds. Science 291: 2598–2600.

